# TargetMITO: A rule-based model for generating highly functional synthetic mitochondrial targeting sequence in yeast

**DOI:** 10.64898/2026.02.22.707306

**Authors:** Kewin Gombeau, Raymond Wan, Joshua S. James, Déborah Tribouillard-Tanvier, Yizhi Cai

**Affiliations:** Manchester Institute of Biotechnology, University of Manchester, 131 Princess Street, M1 7DN, Manchester, United Kingdom; CNRS, IBGC, UMR 5095, University of Bordeaux, F-33000 Bordeaux, France

**Keywords:** mitochondria, synthetic biology, mitochondrial targeting sequence, allotopic expression

## Abstract

Mitochondria are essential organelles containing their own genomes, encoding a few proteins essential for energy production. Most of the mitochondrial proteins are nucleus-encoded, translated as precursors in the cytoplasm, with a large fraction of these precursors properly addressed by an N-terminal mitochondrial targeting sequence (MTS). These MTS share common features but no consensus sequence can explain their functionality nor the precursors-specific determinants of mitochondrial import. To decipher this mechanism, we created a simple computational model to generate highly functional synthetic MTS while maintaining a tight control on the design parameters. Using the budding yeast, we demonstrated the presence of precursors-specific signatures in addressing artificially nucleus-relocated OXPHOS proteins. We also show the ability of six promising candidate synthetic MTS to address a fluorescent reporter to human mitochondria cells. Our research work confirms the uniqueness of the MTS-passenger protein synergy and takes us one step closer towards improving gene therapy-based treatment of mitochondrial diseases.

**Graphical Abstract:** 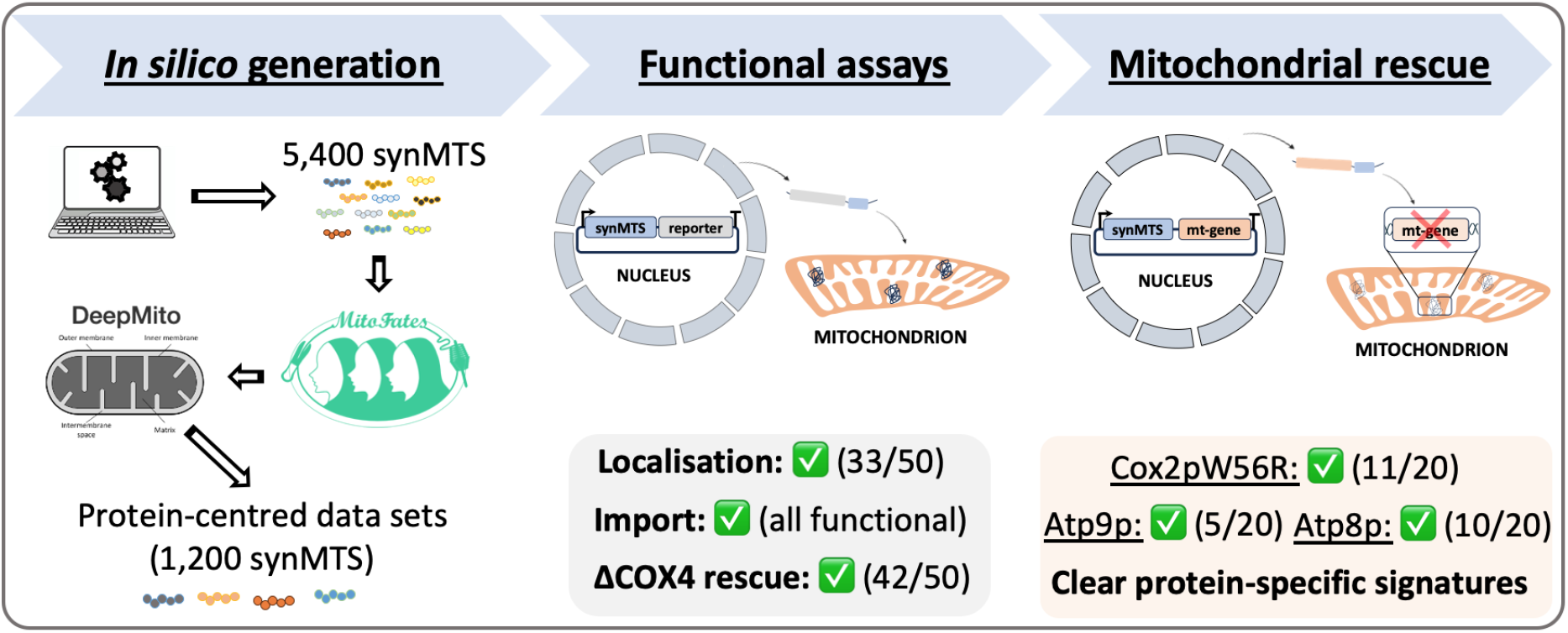

## Introduction

Mitochondria are crucial eukaryotic organelles hosting multiple essential biological processes encompassing the biosynthesis of branched chained amino acids and iron-sulphur clusters, calcium homeostasis, TCA cycle, lipids metabolism, regulation of apoptosis and ATP production through the oxidative phosphorylation system (OXPHOS system) (24). While most of these functions are encoded in genomic DNA, the OXPHOS system of most organisms presents a unique feature: a dual-genetic origin, from both the nuclear and mitochondrial genome (mtDNA) (38). Interestingly, on the one hand, this OXPHOS system involves about 100 different proteins species among which a small fraction (13 in human and 7 in budding yeast) remains encoded in the mtDNA, along with a set of tRNAs and rRNAs essential for the intraorganellar translation of mitochondrial proteins (38). On the other hand, the nucleus-encoded mitochondrial proteins are synthesised as protein precursors in the cytoplasm prior to being targeted to mitochondria. A large fraction of these precursors contains at their N-terminus a cleavable positively charged amphipathic presequence signal (with a length varying between 10 to 120 amino acids), often forming an alpha-helix, and referred to as the mitochondrial targeting sequence (MTS) (12). These MTS play a crucial role in ensuring an efficient mitochondrial import of the protein precursor by guiding it towards its corresponding mitochondrial sub-compartment where it is matured through the action of mitochondrial peptidases excising the MTS (12). For more than 40 years, researchers have explored the determinants of mitochondrial import by trying to identify a potential amino acid pattern explaining the functionality of these MTS (6; 1; 11; 23). The results demonstrated that rather than a specific amino acid consensus sequence, the overall composition, and particularly the balance between hydrophobic and hydrophilic positively charged residues is central in explaining the functionality of the presequences. Recently, artificial presequences were derived from a data set of naturally occurring MTS using a convolutional neural network and proven able to target soluble proteins (i.e. either a GFP or enzymes) to mitochondria (6). In keeping in line with earlier works, their physicochemical properties matched those of natural presequences: amphipathic peptides enriched in arginine with an alpha-helical structure.

While the aforementioned study confirmed that artificial sequences mimicking natural ones can be used to target simple soluble proteins to mitochondria, for more complex cargos, MTS cross-compatibility does not always exist. We have highlighted this issue recently trying to improve the nuclear expression of the artificially relocated *COX2-W56R* gene in *Saccharomyces cerevisiae* (the budding yeast)(15). Also referred to as allotopic expression, the artificial nuclear relocation of mitochondrial genes is widely used to both understand the forces responsible for the retention of mitochondrial protein-coding genes in the mtDNA as well as to create gene therapy-based treatments for mtDNA-linked diseases (3). This method requires: i) recoding the mitochondrial gene to match the nuclear genetic code; ii) attaching to its N-terminus an MTS to properly address the cytoplasm-synthesised protein to the mitochondrial inner membrane; and iii) adjusting the physicochemical properties of the relocated protein to prevent cytoplasmic aggregation and/or mislocalisation (9; 37; 31; 10; 4; 29). In our study on *COX2-W56R* nuclear relocation in yeast, we demonstrated that despite identifying new functional mutations improving Cox2p-W56R import and sorting, a significant fraction of the allotopic protein was not imported into the organelle, regardless of the native MTS used (15). This is in line with previously reported results (8). They tested 20 different individual MTS for their ability to target nine different allotopic proteins in human cells and the results pointed out a low efficiency of the endogenous sequences highlighting the lack of effective MTS inter-compatibility between mitochondrial proteins. This is probably due to both the hydrophobic nature of these transmembrane proteins which impacts their mitochondrial import and the possibility that native MTS have co-evolved with the protein they are attached to, ensuring efficient co-translational import and sorting into mitochondria. As such, creating a simple model to better control the design parameters to generate highly functional synthetic presequences will help decipher the determinants of mitochondrial import. While the common features shared between MTS are well known, exploring the properties conferring MTS a passenger protein specificity will not only help elucidating the mechanism of mitochondrial protein import but aid in the development of new treatments for mtDNA-linked diseases. Indeed, these diseases are among the most common genetic disorders in humans (16), and to date, no effective treatment has been discovered. Multiple routes are being explored, among which gene therapy replacement, involving the allotopic expression of a healthy copy of a mitochondrial gene, is a promising candidate (36; 28; 39).

In this context, we are addressing in the present paper the issue of artificially nucleus-relocated mitochondrial protein targeting to mitochondria. In particular, we decided to shift the paradigm on the study of the determinants of MTS import properties. Usually, the approach chosen to analyse MTS composition and function is performed on data sets of sequences either pooled from a single organism or gathered from multiple life kingdoms. This has been useful in identifying the most commonly shared features between MTS (i.e. Positively charged arginine-rich and amphipathic peptides with an *α*-helical structure carrying a mitochondrial peptidase cleavage site). Unfortunately, a direct consequence of this aggregated analysis is the dilution of the precursors-specific MTS composition that explains the addressing to a specific mitochondrial sub-compartment. This is crucial information, as highlighted in our recent research on improving *COX2-W56R* gene allotopic expression in yeast (15). While looking for potential endogenous MTS capable of efficiently addressing the protein to mitochondria, our findings demonstrated that *OXA1* MTS was the most efficient, even when compared to endogenous MTS addressing nucleus-encoded Complex IV proteins (i.e *COX6, COX11* or *COX15* MTS) (15). This lack of efficient MTS cross-compatibility clearly demonstrates the need to focus on precursors to identify the determinants of their mitochondrial import. This hypothesis was recently corroborated proposing that MTS contain a protein-specific priority code to efficiently address their cargo to its mitochondrial sub-compartment (32).

To identify the passenger protein-specific import determinants, a large set of *in vivo* functionally validated presequences (MTS) is required. Unfortunately, this remains currently infeasible due to the limited availability of public data sets and the poor cross-compatibility of MTS. As such, we generated a simple model to create highly functional, entirely unique and new-to-nature synthetic MTS (synMTS) able to efficiently target artificially nucleus-relocated mitochondrial OXPHOS proteins in the budding yeast. Using known universal MTS’ characteristics, we defined four simple rules to create our sequences: i) a sequence with a length between 40-120 amino acid starting with a methionine; ii) the presence of a mitochondrial peptidase cleavage site at the end of the sequence; iii) the presence of a repetitive pattern of arginine and lysine residues; and iv) a doubled occurrence of arginine/lysine every 30-40 amino acids. Using existing prediction software (MitoFates (13) and DeepMito (34)), we benchmarked our synMTS *in silico* against commonly used MTS in artificial nuclear relocation experiments in yeast (the two gold standards: *OXA1* MTS and *SU9* MTS). Based on the obtained prediction scores, we selected a set of 50 unique sequences whose functionality was assessed *in vivo*. First, we applied various assays starting with a fluorescence-based localisation assay to confirm the ability of the synMTS to efficiently target mitochondria. Using the IQ-Compete assay (32), we measured the import properties of our synthetic presequences. Next, we tested the ability of these synMTS to address the nucleus-encoded Cox4p by virtue of restoring the respiratory growth of the *COX4* deletion strain. Then, we focused on characterising the mitochondrial targeting of three allotopically expressed proteins (Cox2p-W56R, Atp9p and Atp8p) by evaluating the restoration of the respiratory growth of their corresponding deletion strains, measuring the oxygen consumption rates in whole cells, and when possible, quantifying the levels of accumulated allotopic proteins. The results demonstrated that our model could generate highly functional presequences with clear instances of passenger protein-specific functionality. Finally, we selected six of the most promising synMTS and confirmed their ability to route a fluorescent reporter to mitochondria in human cells. This study lays the foundation to our future research work, where we intend to analyse thousands of synthetic presequences in order to identify the passenger protein-specific synMTS’ features, ultimately aiding the development of gene therapy strategies for mitochondrial diseases in humans.

## Methods

### synMTS Generation

Our rationale for the design of our synthetic MTS involved maintaining a repetitive pattern of hydrophilic residues (arginine an lysine) while filling empty intervals with randomly chosen residues excluding the hydrophilic ones. Hydrophylic patterns in this work were calculated using the amino acid hydrophilicity scale of (19) (See also Table S9.)

Based on the hydrophylic patterns of *OXA1* MTS and *SU9* MTS, several methods for synMTS generation were derived. We begin by defining some notation in order to explain the properties of our synMTS. We represent the set of all 20 canonical amino acids as (𝒜). We segment 𝒜 into separate sets as follows. The set ℋ_1_ includes the two strongly hydrophilic and positively charged amino acids (lysine (K), and arginine (R)), while ℋ_2_ includes the next two strongly hydrophilic negatively charged amino acids (aspartic acid (D), glutamic acid (E)). Meanwhile, ℋ is defined as the union of these two sets. In other words, ℋ_1_ ∈ {K, R}; ℋ_2_ ∈ {D, E}; and ℋ = ℋ_1_ ∪ ℋ_2_. Finally, the remaining 16 amino acids are placed in the set *𝒪*.

Using these sets as building blocks, we incrementally apply design rules to guide the creation of our synMTS data sets through nine different methods whose parameters are summarised in Table 1. Each column in this table indicates a feature, which from left-to-right, is associated with an increased stringency in the applied design. Method 1 simply requires the presequence to start with a methionine (M). The remainder of the presequence is calculated randomly from all of the amino acids using a uniform distribution, so that the total presequence is between 40 to 120 amino acids in length. As a regular expression, this is represented as M(*𝒜*)^+^.

**Table 1.**
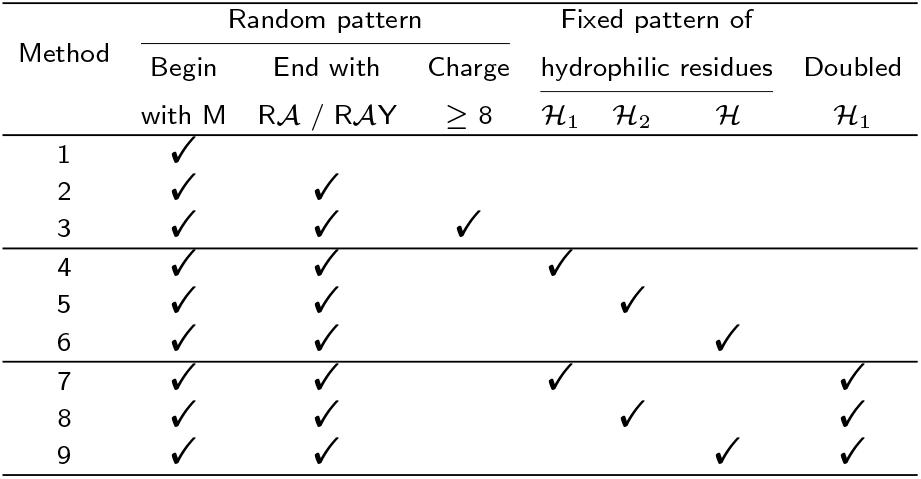
Summary of the nine methods considered by this work for the creation of the synMTS data sets. Each column represents a property that the method has, as indicated by ✓. In this table, ℋ_1_ = lysine and arginine, ℋ_2_ = aspartic acid and glutamic acid, ℋ = ℋ_1_ ⋃ ℋ_2_, and 𝒜 = all the amino acids. M = methionine, R = arginine, Y = tyrosine. Horizontal lines have been added to improve readability.

In addition to the starting methionine requirement, Method 2 stipulates that each MTS must also include a known mitochondrial peptidase cleavage site, corresponding to either an arginine at the −2 position from the C-terminal (R-2) or arginine at the −3 position followed by a tyrosine at the −1 position (R-3 and Y-1) (14). This is represented as M*𝒜* +(R*𝒜*Y^{0,1}^) as a regular expression.

Building on Method 2, Method 3 aims to increase the net positive charge, a crucial feature of MTS. We achieve this by randomly permuting the amino acids in favour of the two *ℋ*_1_ residues until the net positive charge of the entire presequence is eight or higher. This is represented as M*𝒜* ^*∗*^(*ℋ*_1_ +*𝒜* ^*∗*^)^+^(R*𝒜*Y^{0,1}^).

The remaining 6 methods include hydrophilic residues (i.e., *ℋ*_1_, *ℋ*_2_, or *ℋ*, as indicated in Table 1) at intervals of every 3-5 amino acids so as to mimic the observed repetitive pattern seen in the reference MTS (Figure 1(A) and Figure 1(B)). In between these hydrophilic residues are amino acids randomly taken from *𝒪*. Moreover, we observed in the reference MTS hydrophilicity patterns few instances of hydrophilic residues that were doubled. Thus, for the last 3 methods, we decided to explore the influence of this doubled hydrophilic positions by inserting an amino acid from *ℋ*_1_ in the first 30–40 amino acids. For synMTS longer than 40 amino acids, these intervals are used: [30, 45]; [60, 75]; [90, 110]. A summary of the different methods deployed in this study is provided in Table 1. Regular expressions that describe these nine methods are provided in Table S10.

**Figure 1.**
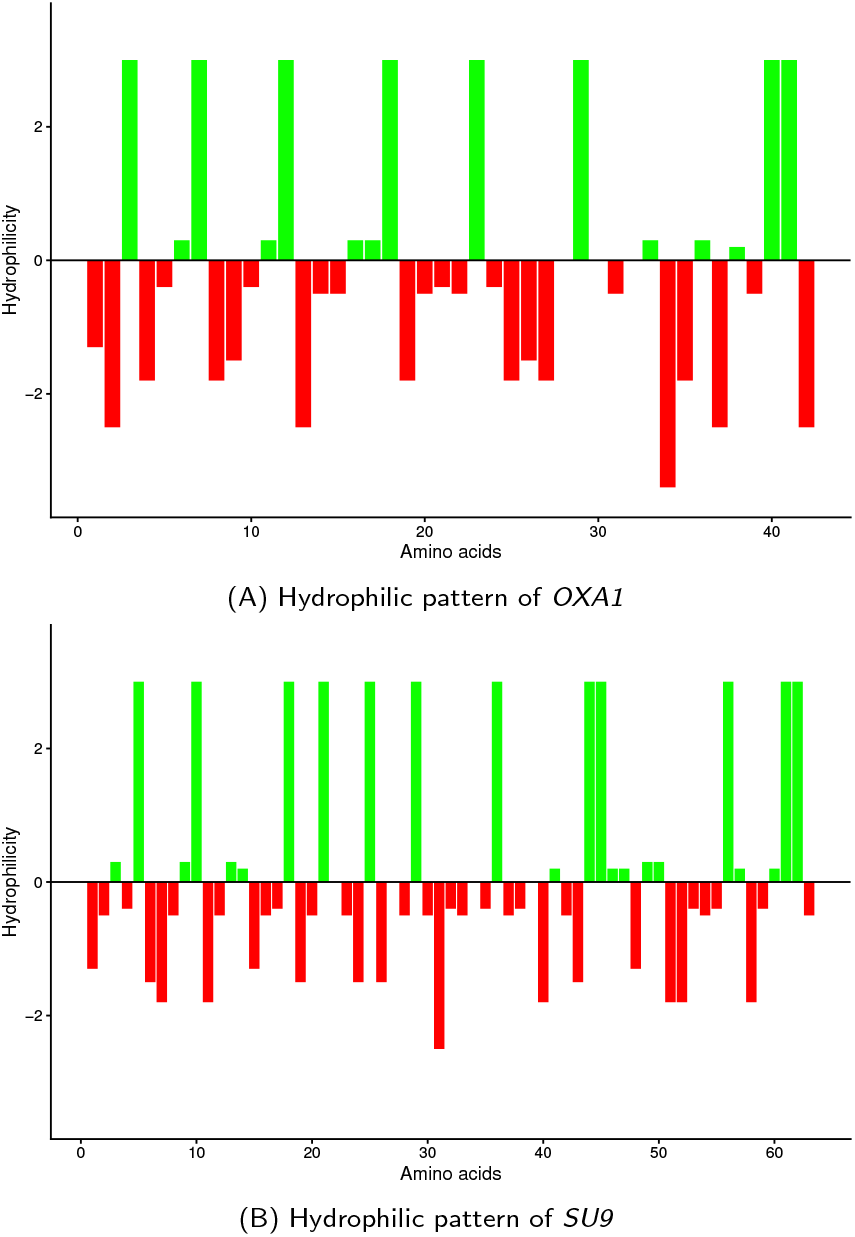
Derivation of the methods considered by this work for generating synMTS. Hydrophilic patterns of the reference (A) *OXA1* and (B) *SU9* MTS used in artificial nuclear relocation experiments. The large green bars represent the hydrophilicity value of either arginine or lysine.

We confirmed that their lengths (Figure S2) and total net charges (Figure S3) were consistent with the set of rules imposed across the different methods.

All computational experiments in this work were performed on a Intel^®^ Core^™^ i9-14900K CPU with 32 GiB RAM running Ubuntu 25.04 . The software consists of a series of scripts that were executed with Perl version 5.32.1 and R version 4.4.2. These scripts have been combined into a Snakemake version 8.26.0 workflow (21).

### Homology search with BLASTP

Homology search of the generated synMTS data sets using BLASTP was performed to validate the uniqueness of the generated sequences. The RefSeq protein database downloaded from https://ftp.ncbi.nlm.nih.gov/blast/db/ on 28 August 2025 was used (2).

BLASTP version 2.16.0+ was set up by installing the Ubuntu package “ncbi-blast+” version 2.16.0+ds-6build1. The synMTS data set was queried using the following command:

~~~
time blastp −query . / query . fasta −db \
refseq_protein −out results . tab −evalue 1 e−5 \
−outfmt 7 −num threads 24
~~~

Here, an E-value cut-off of 1 × 10^−5^ and 24 threads of execution was employed. On the computer used for our experiments, this took almost 28 hours to run on our set of 5,400 synMTS sequences.

### Benchmarking the synMTS with MitoFates

MitoFates (13) was used to evaluate the synthetic MTS produced. Since MitoFates had been trained on *S. cerevisiae* data sets of experimentally confirmed mitochondrial proteins, we considered it as an adequate tool to benchmark the different methods used to generate our synthetic MTS.

As expected, we observed a distinct variation in the obtained presequence probability scores across the different methods which depended on the stringency of the applied rules (Figure S4). The data set of random sequences starting with a methionine (Method 1) returned low probability scores, which was not improved by the addition of known mitochondrial cleavage sites (Method 2). While a slight improvement was observed when the synthetic MTS had to carry a minimum of eight positive net charges (Method 3), the most substantial enhancement was achieved by enforcing a regular pattern of of positively charged residues (Methods 4 and 7). Conversely, incorporating a mixture of hydrophilic charged residues (Methods 6 and 9) led to a significant reduction in prediction scores, while designs utilising only negatively charged residues completely eliminated any positive predictions (Methods 5 and 8), confirming the need to only use positively charged hydrophilic residues.

To appraise the dependency of the calculated MitoFates probability scores on the protein sequence appended to the different MTS, we included the negative control Hac1p (a randomly selected nuclear transcription factor). The obtained results confirmed that MitoFates predictions are robustly consistent across the different conditions regardless of the appended protein (Figure S4).

To further confirm these findings, for each protein, we merged the data sets from Methods 4 and 7 into a single data set, as depicted in Figure S1(B). We then sorted the scores from highest to lowest and took the top N scores, for various values of N. Finally, we intersected the lists for every possible pair of mitochondrial proteins (Figure S5 shows these results). For example, for the pair Atp8p and Atp9p, 92.8% of the top 250 scores for these two passenger proteins overlap (the dotted red vertical line in this figure). As this graph confirms, the results are highly similar across the different conditions and rather insensitive to the appended protein.

Our analysis with MitoFates was performed using the web server at the following address: https://mitf.cbrc.pj.aist.go.jp/MitoFates/cgi-bin/top.cgi.

### Exploring the “synMTS-passenger protein” synergy with DeepMito

DeepMito employs a convolutional neural network to classify proteins and predict sub-mitochondrial localisation (34). When using DeepMito through the web server, TPpred3 and BaCelLo are also called (30; 20; 35) and provide a binary answer as to whether or not the protein in question is a mitochondrial protein.

Analysis with DeepMito was performed using the web server at the following address: https://busca.biocomp.unibo.it/deepmito/, with the prediction scores obtained for the selected synMTS given in Table S5.

### Yeast culture and transformation

We used the previously described biolistic transformation procedure and the *ARG8*^*m*^ marker to generate the different mitochondrial deletions (5). Briefly, the different deletion cassettes were delivered in the rho0 DFS160 yeast strain which was then crossed with the wild type strain YCy5051 (referred to as WT) to create the different single deletion strains (either YCy6163 (referred to as Δ*atp9::ARG8*^*m*^), or YCy6976 (referred to as Δ*atp8::ARG8*^*m*^). The W303 and *COX4* deletion strains are kind gifts from Dr. Natalie Niemi. All liquid precultures were performed in liquid synthetic media with corresponding dropouts, starting from fresh transformants and incubated without shaking. We performed the spot test assays on rich media (1% yeast extract and 2% peptone) supplemented with either 2% glucose or 2% glycerol. We spotted, for each strain, a 10-fold serial dilution, starting from a concentration of 1 OD_600nm_ unit/mL and growing the cells at 30°C for the indicated time. A list of the yeast strains used and generated in this study is provided in Table S1.

### Plasmids assembly strategy

In the present study, the different transcriptional units were assembled using the golden gate-based method called YeastFab (18); with modified overhangs to enable the incorporation of one or two MTS between a given promoter and a coding gene, as previously reported (15). The golden gate reactions were transformed in TOP10 *Escherichia coli* competent cells and the expression vectors purified using the E.Z.N.A plasmid miniprep kit (Omega Bio-tek, Norcross, GA, USA). Then, these expression vectors were transformed in yeast using the classic LiOAc-based transformation protocol isolating the correct transformants on synthetic solid media with corresponding amino acid dropouts. Mammalian synMTS-mKate2 expression vectors were assembled using the EMMA Toolkit and a modified EMMA receiver vector backbone hosting a Cox8 MTS-mNeonGreen fluorescent marker control cassette (25). Lists of the primers (Table S2) and plasmids (Table S3) are provided in the supplementary files.

### Fluorescence-based procedures

Using fluorescence microscopy, we appraised the capability of our synthesised synMTS to address a Cherry fluorescent protein to mitochondria. To achieve this, we imaged cells during their exponential phase of growth using an inverted epifluorescence microscope (Nikon Eclipse TE2000U) equipped with an 100× immersion objective and a standard FITC filter. The results are presented in Figure 2. The fluorescence analyses conducted on human cells were conducted using a Cytation5 (Agilent) plate reader equipped with and inverted microscope, a 40x objective and the corresponding filters (Figure 10).

**Figure 2.**
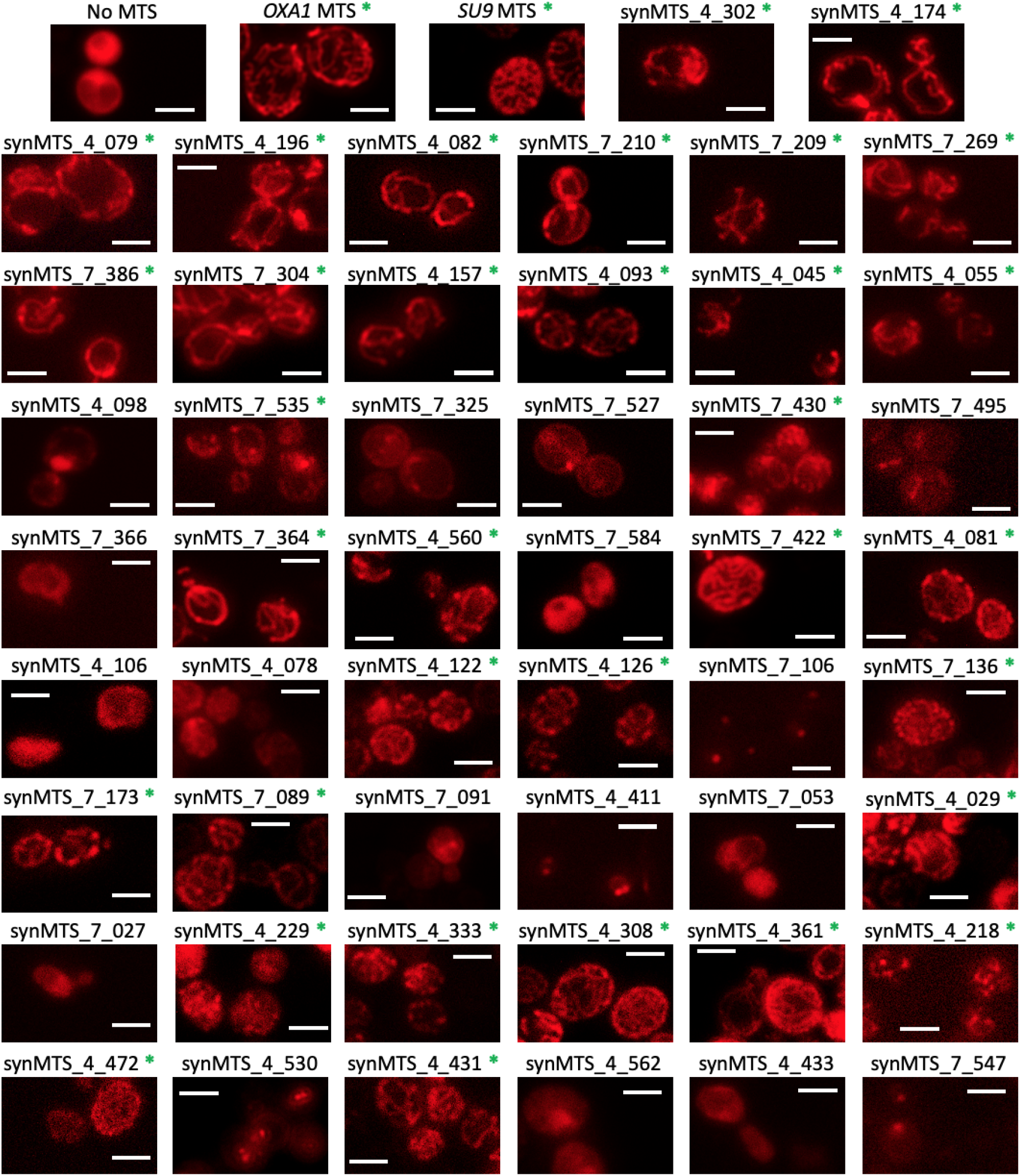
Fluorescence imaging of the BY4741 rho+ strain expressing the different MTS-Cherry constructs. Scale bar = 5 *µ*m. The green asterisks denote a satisfactory addressing conferred by the corresponding synMTS.

To quantitatively measure the import properties conferred by our synMTS, we used the IQ-Compete assay (32), adapting the procedure to make it compatible with our YeastFab assembly method (18). Briefly, the Tobacco Etch virus protease (uTev) was combined with the different synMTS and co-expressed, along with the GFP-quencher-degron, on two different low-copy vectors (see Table S3 for a list of plasmids generated). The GFP signal obtained under each condition was normalised to the cell density in each well and expressed as a percentage of the signal measured in the condition where uTev is cytoplasmic (no MTS added) (Figure 3. The transformants were grown until saturation without shaking for 24h. Then the precultures were transferred in synthetic media with the corresponding dropouts supplemented with 2% glucose and the fluorescence monitored for 24h using a Cytation 5 plate reader (Agilent) and normalised, for each well, to cell density. The GFP fluorescence signal was used when cells were in their exponential phase of growth. The results are presented in Figure 3 and the corresponding data is presented in Table S6.

**Figure 3.**
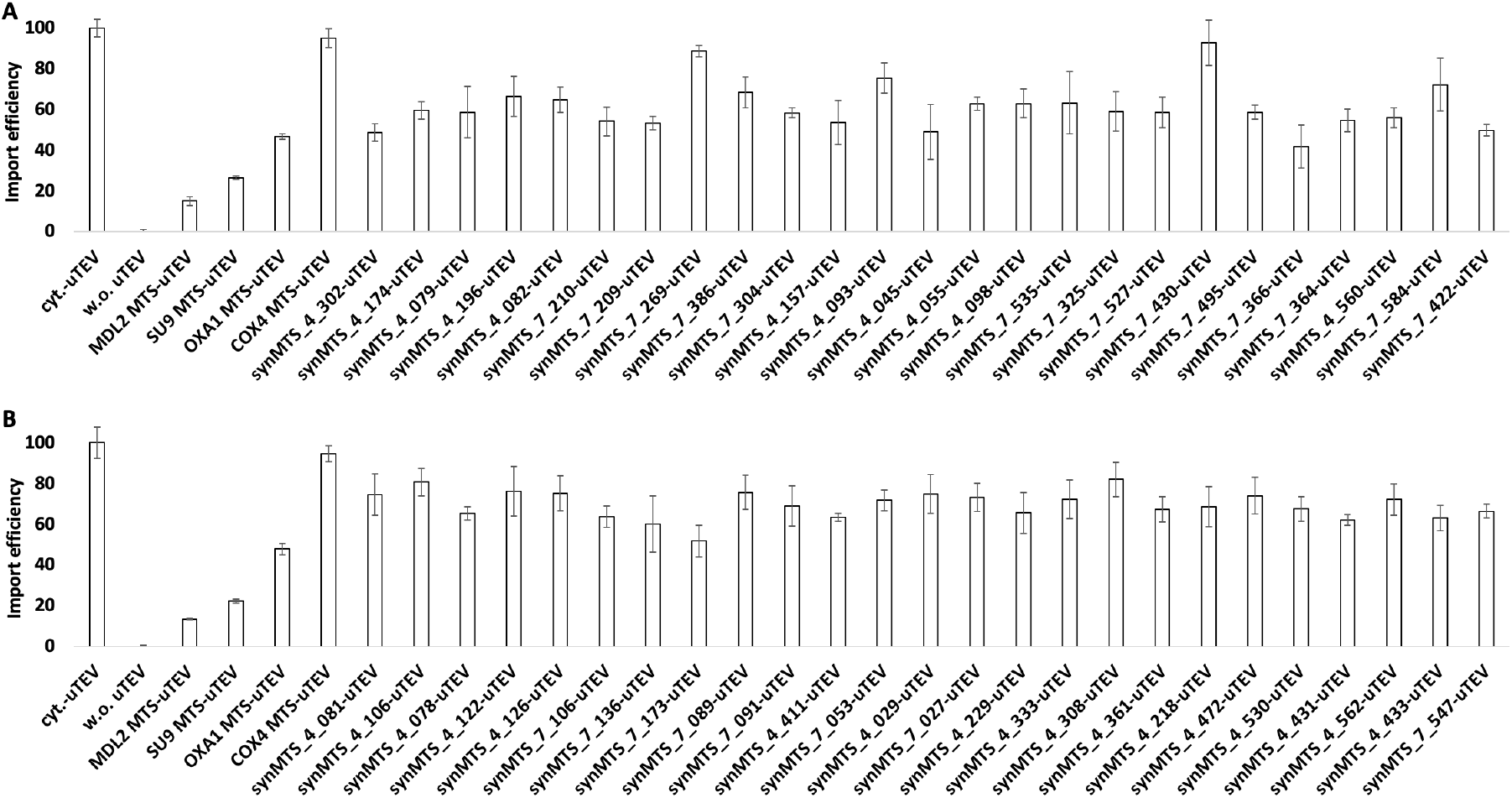
Results of the IQ-Compete assay performed using the reference MTS and the 50 unique synthetic presequences generated in this study. The data presented corresponds, for each condition, to the mean +/-SD of the GFP signal measured for three biological replicates normalised to cell density in each well and expressed as a percentage of the signal measured in the condition “cyt.-uTEV”.

### Biochemical analyses of the respiratory metabolism

We quantified the ability of the selected allotopic expression systems to complement the deletion of their corresponding mitochondrial gene (either *COX2-W56R, ATP9* or *ATP8* ) by measuring both the rates of oxygen consumption (OCR, using a Clark electrode from Hansatech, UK) and the level of allotopic protein accumulation. For each of the different allotopic strains, freshly obtained transformants were inoculated in static synthetic liquid media (with dropouts corresponding to the auxotrophy markers used and supplemented with 2% glucose) and grown overnight at 30°C. The next day, the cultures were transferred to 6 mL of rich medium supplemented with 2% glycerol (YPGly 2%) at a starting OD_600nm_ of 0.25/mL and grown overnight at 30°C with shaking. Finally, the cells were harvested during their exponential growth phase to measure the OCR on whole-cells or to prepare total protein extracts. The raw data of OCR measurements on whole cells are available in Table S7.

The protein extracts were subjected to SDS-PAGE (22) and Western Blotting. The abundance of Cox2p was measured using a commercially available primary antibody (Abcam, ref ab110271, dilution 1:2500). Since no antibodies are available for the immunodetection of the yeast Atp9p and Atp8p, we created C-terminal FLAG-tagged strains. Unfortunately, this was either creating a growth defect, or leading to inconsistent protein detection. For Atp9p, we also tested an *in-house*-generated polyclonal antibody, but the abundance of the protein was too low in our allotopic strains and no exploitable signal could be detected (data not shown). The Cox2p detection signal was normalised, in each condition, to that of Pgk1p (Abcam, ref ab113687, dilution 1:5000) and expressed as a percentage of the WT level. The raw data of Western Blot analyses are available in Table S8.

### Mammalian cell culture and transfection

HAP1 cells (Horizon Discovery) were cultured in Iscove’s Modified Dulbecco’s Medium with Glutamax supplement (Gibco) and 10% FBS in a humidified incubator at 37°C and 5% CO_2_. One day prior to transfection, 100,000 cells were seeded into 12-well plates. Lipofectamine 3000 (2µL P3000, 3µL L3000) was used to transfect 1 µg of plasmids DNA per well. Cells were cultured for 24 hours prior to imaging.

### Statistical analyses

All experiments were conducted with three independent biological replicates per condition. Prior to group comparisons, data were assessed for normality and homogeneity of variance using the Shapiro–Wilk and Levene tests (*α* = 0.05). When both assumptions were met, a parametric one-way ANOVA with Dunnett’s post-hoc correction was applied; otherwise, a Kruskal–Wallis test followed by Mann–Whitney comparisons (*α* = 0.05) was used. Statistical analyses were performed in RStudio (version 4.2.3), and reported *p*-values are corrected values.

### Software and data availability

Source code for generating and evaluating the synMTS data sets is available through this GitHub repository https://github.com/the-cai-lab/synMTS.

The generated synMTS data set of 5,400 synthetic MTS sequences and our results from submitting these sequences to MitoFates and DeepMito are also included within this GitHub repository.

## Results

### Design and cross-validation of novel synthetic MTS

The design of our synMTS relies on the examination of the physicochemical properties of the *OXA1* and *SU9* MTS which provided the best results in addressing Cox2p-W56R and Atp8p/Atp9p to mitochondria, respectively, in previous artificial nuclear relocation experiments in yeast (26; 37; 4; 15). The hydrophylic patterns of *OXA1* MTS (8 positive net charges) and the *SU9* MTS (12 positive net charges) are shown in Figure 1(A) and Figure 1(B). These two examples illustrate the features that are essential in determining the functionality of MTS: i) a total positive net charge; ii) an amphipathic alpha-helical structure; and iii) a frequent occurrence of arginine and lysine residues.

Using this knowledge as our foundation, we developed different methods for generating the synMTS *in silico*. The procedure we opted for is detailed in the Methods section and resulted in the development of nine different methods (Table 1C). For each of these methods, three independent replicates of 200 synthetic MTS were produced (600 presequences per method), yielding 5,400 synMTS in total (as depicted in Figure S1(A)). A homology search for all of these sequences using BLASTP against the RefSeq protein database returned no hits, confirming that the generated sequences are entirely new-to-nature.

Next, our aim is to identify which methods yielded the best synthetic MTS. Here, we turned to the MitoFates web server, a prediction software that analyses the N-terminus region of the submitted protein sequence and returns a presequence probability score indicating the likelihood of the presence of an MTS.

MitoFates requires the presequence to be fused with a passenger protein as input. We appended four proteins to these presequences in order to use these tools to explore the synMTS-passenger protein synergistic relationship. Three of these are mitochondrial proteins localising to the inner mitochondrial membrane (Cox2p-W56R, Atp8p and Atp9p) previously artificially expressed from the nucleus using either *OXA1* or *SU9* MTS (26; 37; 4; 15). The fourth, a negative control, is a nuclear transcription factor (Hac1p) for assessing the robustness of the passenger protein-based predictions.

The prediction scores obtained (Figure S4) revealed that Methods 4 and 7 returned the highest probability scores, regardless of the appended protein, confirming the MTS-centred architecture of the MitoFates analyses. We merged data sets from Methods 4 and 7 (see Figure S1(B)) and performed a pairwise comparison between data sets of synMTS + passenger proteins (Figure S5). These results reinforce the idea that the scores are not affected by the fused passenger protein.

### Exploring synMTS–passenger protein synergy by predicting the sub-mitochondrial localisation

To explore the synergy between our synMTS and the appended proteins, we continued to use the merged data sets from Methods 4 and 7 (see Figure S1(B)) and submitted, for each appended protein, the merged data sets to the DeepMito web server (34). This software provides predictions on: i) whether a submitted protein, analysed in its entirety (MTS + passenger protein), is localising to mitochondria; and ii) the likelihood of mitochondrial sub-localisation using artificial neural networks trained on data sets of mitochondrial proteins (covering multiple kingdoms of living organisms) whose sub-localisation have been experimentally validated. For the former, DeepMito leverages BaCelLo and TPpred3 to return a “Yes/No” prediction of mitochondrial affiliation (30; 20; 35). For the latter, DeepMito provides a score from 0 to 1 to represent the likelihood.

The prediction results obtained confirmed the mitochondrial localisation of more than 90% of the synMTS-mitochondrial proteins precursors, but not for our negative control, synMTS-Hac1 (Figure S6). Regarding the mitochondrial sub-compartment predictions, almost all the predictions yielded a correct affiliation to the inner mitochondrial membrane (higher than 99% for each mitochondrial protein data set, Table S11).

Interestingly, the top N sorted probability scores presented in Figure S7 reveals the presence of a possible passenger protein-specific synMTS ranking for each data set. Among the 100 synMTS with the highest scores across each data set, a significant protein-specific signature in the ranking could be observed with overlaps ranging from 20% to 58% for any two of the three data sets. However, the similarity in the ranking rapidly increases with the number of cumulated synMTS sequences, ranging between 75% to 85% in the top 500 synMTS of each of the 3 lists (dotted vertical line, Figure S7).

### Analysis of the mitochondrial import properties of the selected synMTS

After obtaining these three protein-specific rankings of our synMTS data set (Figure S1(C)), we explored the functionality of a subset of the ranked sequences *in vivo*. To select the sequences to be tested, we opted for a hypothesis-driven selection rather than a random selection of sequences. Indeed, as the core hypothesis of this study is to create new-to-nature and highly functional synMTS to be used in nuclear relocation experiments, we selected sequences based on their obtained DeepMito ranking hoping to isolate the best ones. We picked the synMTS with ranks of 1 to 5, 398 to 402, 798 to 802 and 1196 to 1200 (Figure S1(C)). Among the 60 sequences shortlisted across the three data sets, 50 were unique (these sequences are provided in Table S4). Using SnapGene^®^ version 4.1.9 (17), we generated the corresponding yeast codon optimised DNA sequences (Table S4) (27) which were then chemically synthesised by Twist Bioscience (South San Francisco, CA, USA).

First, we tested their ability to address to mitochondria a simple non-chemically challenging fluorescent reporter (Figure 2). Using YeastFab assembly (18), we built the different expression vectors, transformed it into our BY4741 rho^+^ strain and performed microscopy fluorescence observations. When the reporter was not associated with an MTS (“No MTS”), the fluorescence signal was present in the entire cell. When the reporter was associated with the reference MTS (“*OXA1* MTS” or “*SU9* MTS”), the signal was only detectable in the reticulated network of mitochondria. When associated with our synMTS, the results obtained revealed that a majority (33/50, or more than 60%) was highly functional and able to efficiently address the reporter to mitochondria (i.e. “synMTS_7_422” or “synMTS_4_361”). Of note, we observed a decreased efficiency in addressing the reporter to mitochondria with decreasing DeepMito scores (Table S5).

The import properties of our synMTS were then further characterised using the IQ-Compete assay (an *in vivo* assay returning an import efficiency inversely correlated to the GFP signal measured)(32). As compared to the original method, encoding the transcriptional units on low copy plasmids instead of a single genomic copy had a minor effect, with slightly decreased import efficiencies for the reference MTS (*MDL2, SU9, OXA1* and *COX4* (32)). Regarding the values measured for our synMTS, the results demonstrated that most of the sequences efficiently imported the uTev protease into mitochondria, with an average import efficiency of 65% (Figure 3, see Table S6 for the percentage values).

Finally, the last test performed involved assessing the capability of our presequences to target Cox4p to mitochondria in a *COX4* deletion mutant strain. Using YeastFab (18), we built the different expression vectors and transformed it into the *COX4* deletion strain. Their performance was evaluated based on their ability to restore the respiratory growth of the *COX4* deletion mutant (see Figure 4). When we expressed the *COX4* ORF with its native presequence, the respiratory growth of the *COX4* deletion mutant could be restored. However, this did not happen when the *COX4* MTS (condition referred to as “No MTS”) was removed. Using the *OXA1* and *SU9* MTS also enabled the restoration of respiratory growth of the deletion strain. When testing our 50 synMTS, we observed that more than 80% (41/50, Figure 4) were also able to target Cox4p to mitochondria, confirming the correctness of the rules used to design our synthetic presequences.

**Figure 4.**
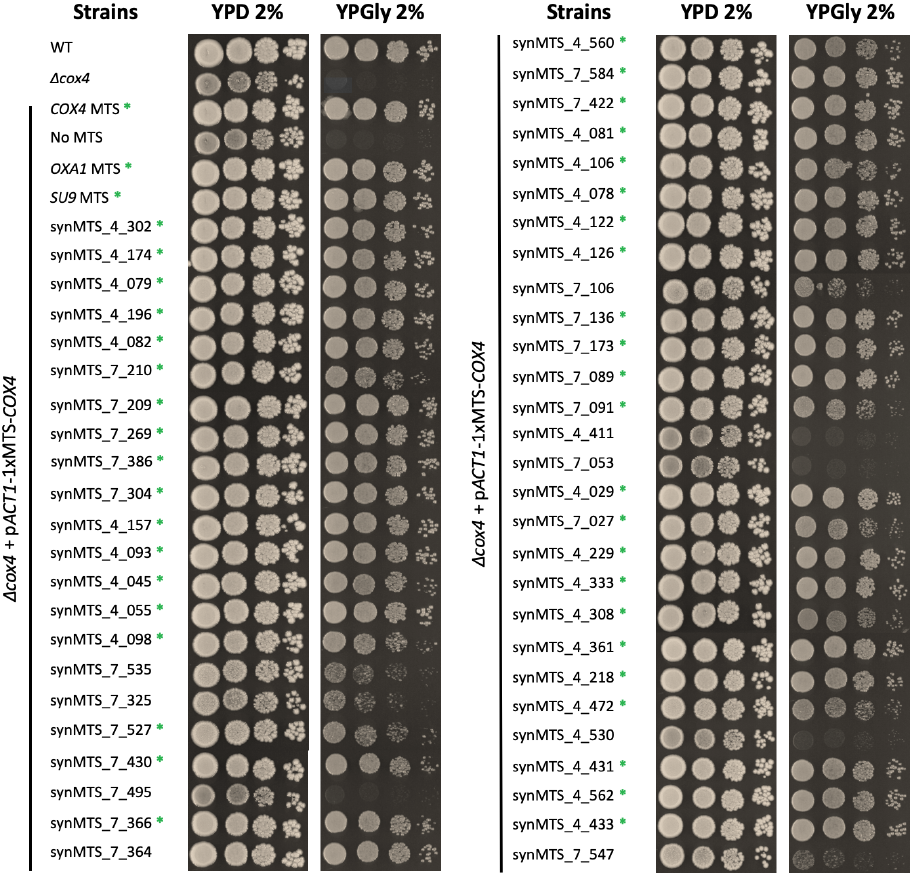
Spot test presenting the growth benefit conferred by using reference or synthetic MTS to address Cox4p to mitochondria in the COX4 deletion strain. The pictures were taken after incubating the plates for three days at 30°C. The green asterisks denote a satisfactory addressing conferred by the corresponding synMTS.

### Functional validation of the selected synMTS

We then tested the synMTS ability to promote the mitochondrial import of three artificially nucleus-relocated allotopic proteins (Cox2p-W56R, Atp9p or Atp8p). The different expression vectors were assembled using the YeastFab assembly (18) with the adaptations previously described (15). Based on our previous observations (15), the *COX2-W56R* gene was encoded on a low-copy vector with one MTS copy under the control of the *ICL1* promoter. These expression conditions offer the best respiratory growth dynamics to identify the improvements conferred by the synMTS (15). The constructs were then delivered in our *COX2* deletion strain and tested for their ability to restore the deletion mutant’s respiratory growth (Figure 5). Regarding *ATP8* and *ATP9* expression conditions, our preliminary tests (data not shown) suggest the use of a multi-copy vector and the *ACT1* promoter. Concerning the MTS copy number, decent growth was observed when Atp8p was associated with one MTS copy. In contrast, Atp9p required two MTS copies. The spot-test assays performed in the *ATP9* and *ATP8* deletion strains are presented in Figure 6 and Figure 7, respectively.

The growth assays revealed that the ability of the tested synMTS-containing expression vectors to complement the respiratory growth of the different mitochondrial deletion strains drastically varied. Indeed, for both Cox2p-W56R and Atp8p, efficient synMTS were observed throughout the entire subsets of tested sequences (Figure 5 and Figure 7). On the other hand, for Atp9p, only the 5 highest-ranking synMTS were able to partially restore the respiratory growth of the mutant (Figure 6(A)). Moreover, for Cox2p-W56R and Atp8p, none of the tested synMTS outperformed the reference MTS sequence (either from *OXA1* or *SU9* ). In contrast, fewer functional sequences were identified for Atp9p, but they markedly improved the *ATP9* deletion mutant’s respiratory growth, even though the impact on the oxygen consumption rates (OCR) measured on whole cells was indiscernible (Figure 6(E)).

**Figure 5.**
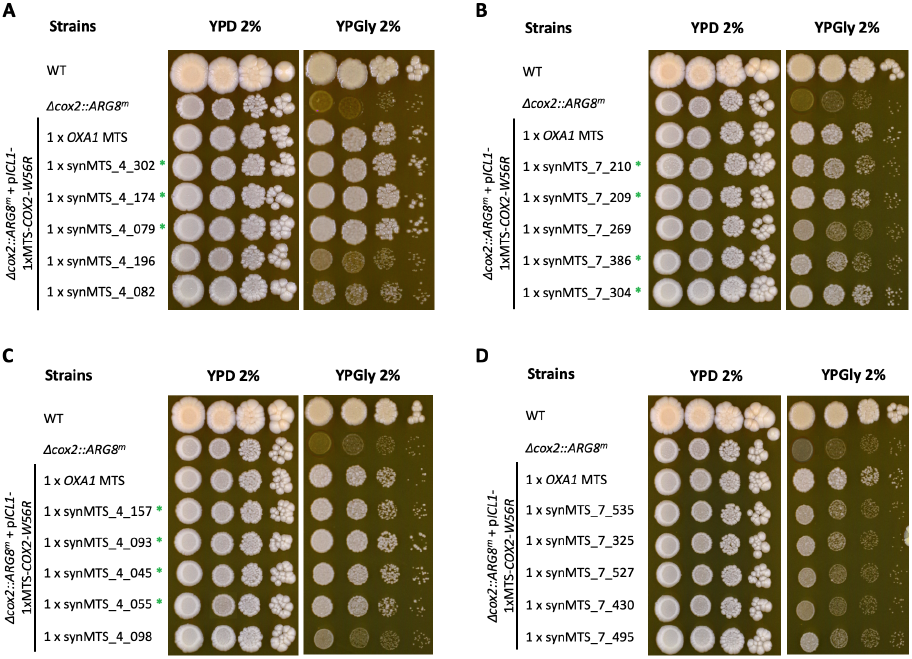
Spot-tests presenting the respiratory growth benefit conferred by the different sets of synthetic MTS in the *COX2-W56R* deletion strain. The sequences were ranked based on their DeepMito scores and sampled at different positions throughout the data set: (A) from 1 to 5, (B) from 398 to 402, (C) from 798 to 802 and (D) from 1196 to 1200. The pictures were taken after incubating the plates for six days at 30°C. The green asterisks denote a satisfactory addressing conferred by the corresponding synMTS.

**Figure 6.**
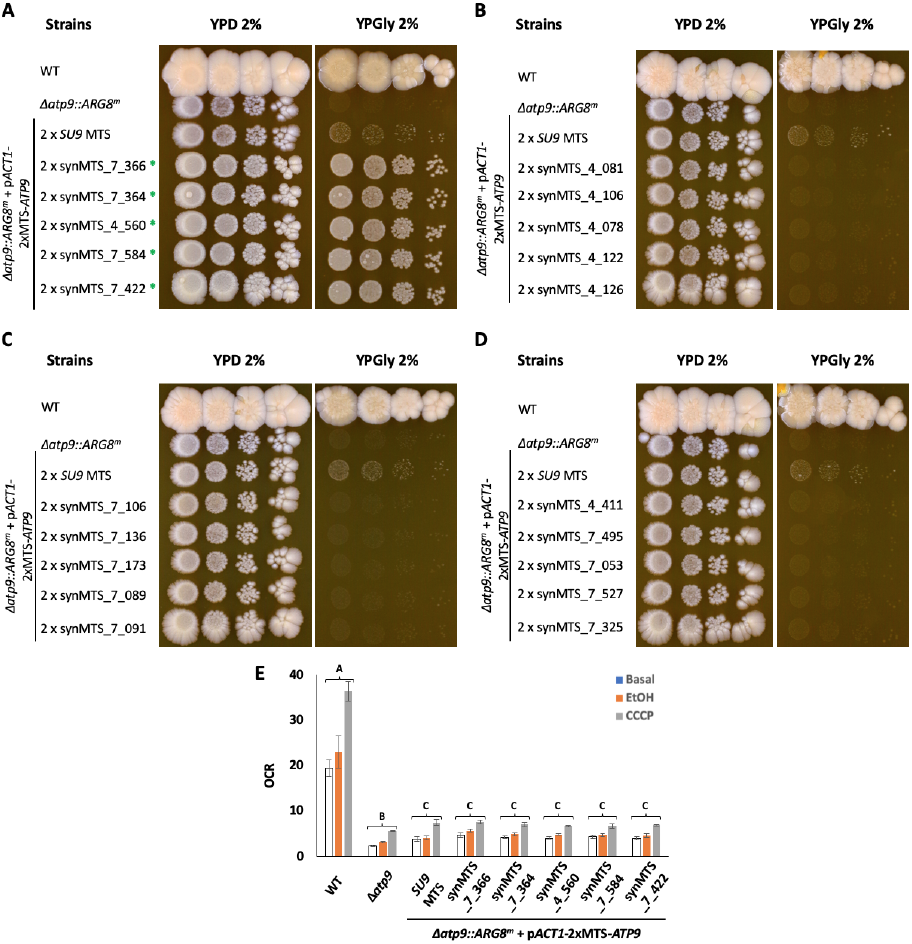
Spot-tests presenting the respiratory growth benefit conferred by the different sets of synthetic MTS in the *ATP9* deletion strain. The sequences were ranked based on their DeepMito scores and sampled at different position throughout the data set: (A) from 1 to 5, (B) from 398 to 402, (C) from 798 to 802 and (D) from 1196 to 1200. The pictures were taken after incubating the plates for ten days at 30°C. We isolated the five best synthetic MTS and measured the corresponding oxygen consumption rates (OCR, nmol.min^−1^.OD_600*nm*_unit^−1^) on whole cells (E). A, B and C: Denotes a statistical difference between conditions (*p* < 0.05). U: Unprocessed protein; M: Mature protein. The green asterisks denote a satisfactory addressing conferred by the corresponding synMTS.

**Figure 7.**
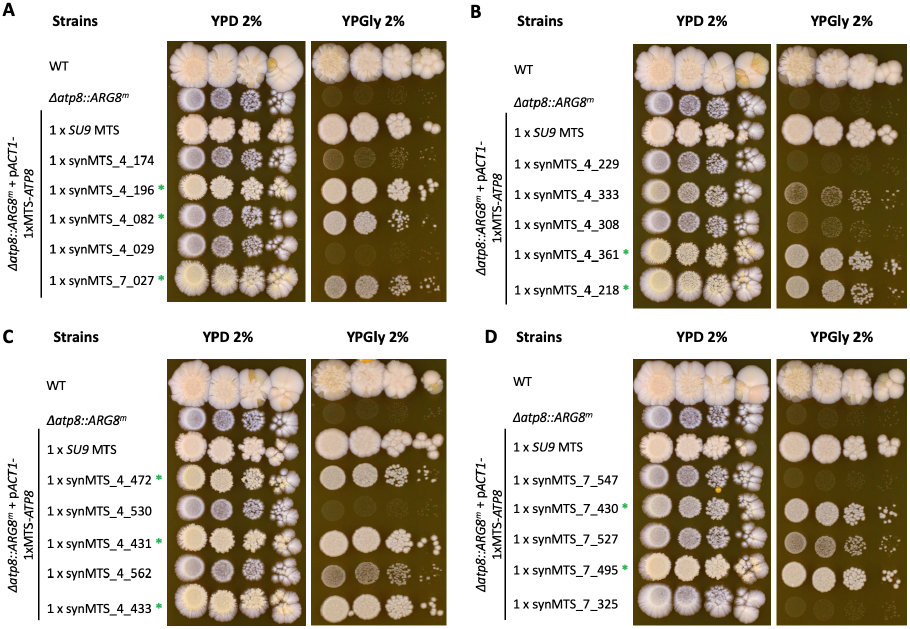
Spot-tests presenting the respiratory growth benefit conferred by the different sets of synthetic MTS in the *ATP8* deletion strain. The sequences were ranked based on their DeepMito scores and sampled at different position throughout the data set: (A) from 1 to 5, (B) from 398 to 402, (C) from 798 to 802 and (D) from 1196 to 1200. The pictures were taken after incubating the plates for ten days at 30°C. The green asterisks denote a satisfactory addressing conferred by the corresponding synMTS.

For both Cox2p-W56R and Atp8p, we repeated the growth experiment shortlisting the most efficient synMTS from this first round of spot-tests and taking pictures at shorter intervals. As only four potential candidate synMTS were identified for Atp8p, we also included the top 5 from Atp9p. Since both proteins belong to the F1/FO ATP synthase complex, we hypothesised that a potential synMTS cross-compatibility could exist. The results obtained for Cox2p-W56R and Atp8p are presented in Figure 8 and Figure 9, respectively.

**Figure 8.**
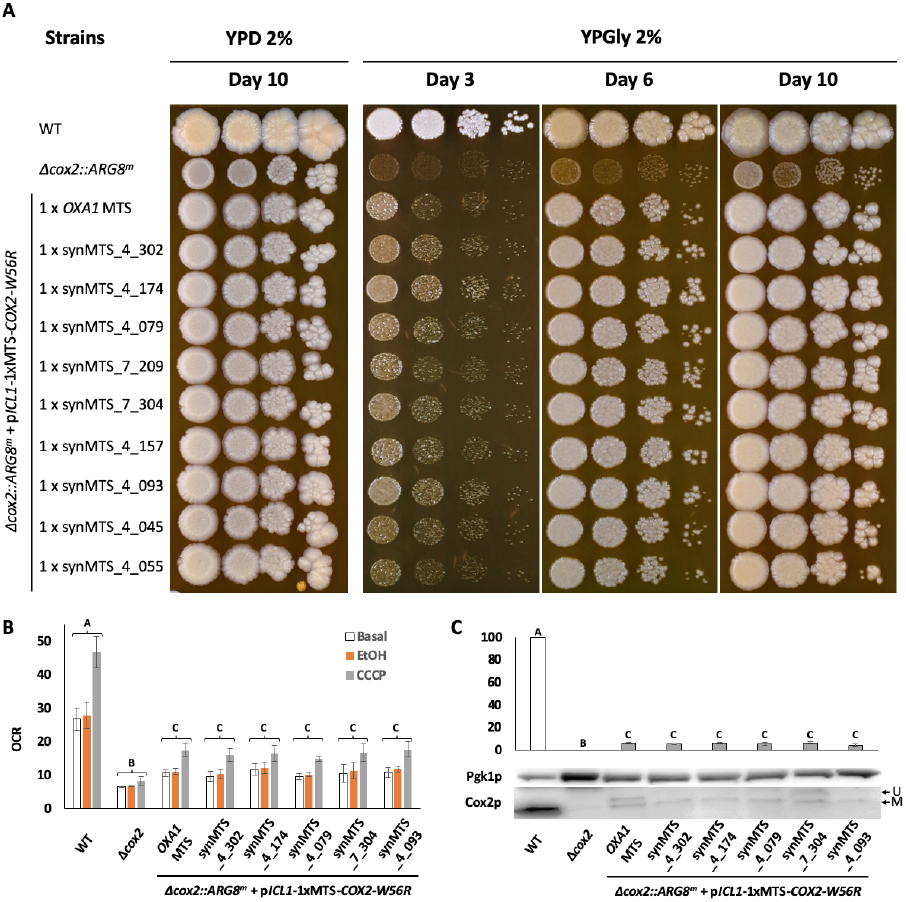
Spot-test confirming the respiratory growth benefit conferred by a subset of nine synthetic MTS in the *COX2-W56R* deletion strain. The pictures were taken after incubating the plates for three, six and ten days at 30°C (A) Then, we isolated the five best synthetic MTS, and measured both the corresponding oxygen consumption rates (OCR, nmol.min^−1^.OD_600*nm*_unit^−1^) on whole cells (B) as well as the level of accumulated Cox2p-W56R (C) A, B and C: Denotes a statistical difference between conditions (*p* < 0.05). U: Unprocessed protein; M: Mature protein.

**Figure 9.**
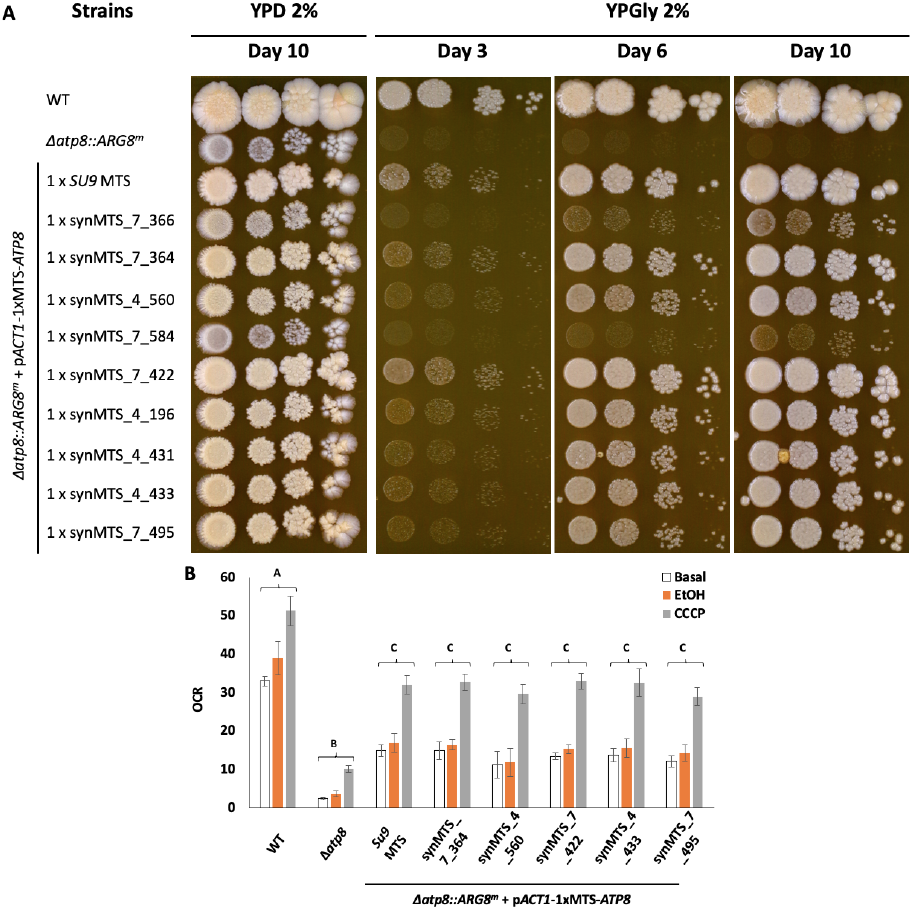
Spot-test confirming the respiratory growth benefit conferred by a subset of nine synthetic MTS in the ATP8 deletion strain. (A) The pictures were taken after incubating the plates for three, six and ten days at 30°C. (B) Then, we isolated the five best synthetic MTS, and measured the corresponding oxygen consumption rates (OCR, nmol.min^−1^.OD_600*nm*_unit^−1^). A, B and C: Denotes a statistical difference between conditions (*p <* 0.05).

For Cox2p-W56R, after 3 days of incubation we identified five synMTS (synMTS_4_302, synMTS_4_174, synMTS_4_079, synMTS_7_304, synMTS_4_093) that appeared as efficient as the *OXA1* MTS (Figure 8(A)). We measured the levels of OCRs on whole cells and the mature Cox2p-W56R abundance, and the results were similar between these conditions, confirming the efficiency of the shortlisted sequences in addressing Cox2p-W56R to mitochondria (Figure 8(B) and Figure 8(C)).

For Atp8p (Figure 9), this second round of respiratory growth analysis revealed that: i) none of the shortlisted sequences from the first set of spot-tests were as efficient as *SU9* MTS; and ii) a partial synMTS cross-compatibility exists between Atp9p and Atp8p. More precisely, among the top 5 sequences from the Atp9p analyses, three were compatible with Atp8p (synMTS_7_364, synMTS_4_560 and synMTS_7_422, Figure 9(A)). The measurement of OCRs in whole cells showed no significant differences compared to the condition using the *SU9* MTS.

### Validation of a subset of synMTS in human cells

Since MTS properties are strongly conserved between yeast and human (7), we hypothesised that any of the functional and passenger protein-specific synMTS characterised in yeast could be a strong candidate for *in vivo* validation in human cell models. We thus tested whether a subset of six selected synMTS (synMTS_4_174, synMTS_7_304, synMTS_4_093, synMTS_7_364, synMTS_7_422 and synMTS_4_196) could also direct the fluorescent mKate2 reporter to mitochondria in human HAP1 cells (Figure 10). To this, we built mammalian expression vectors using the EMMA toolkit ((25)) including a *COX8* MTS-mNeonGreen fluorescent reporter as a positive control for mitochondria localisation and associated the mKate2 reporter with either the reference or candidate synMTS and delivered these constructs in HAP1 cells. The fluorescence microscopy-based co-localisation assay demonstrated that all of the candidate synMTS were able to efficiently route mKate2 to mitochondria (Figure 10).

**Figure 10.**
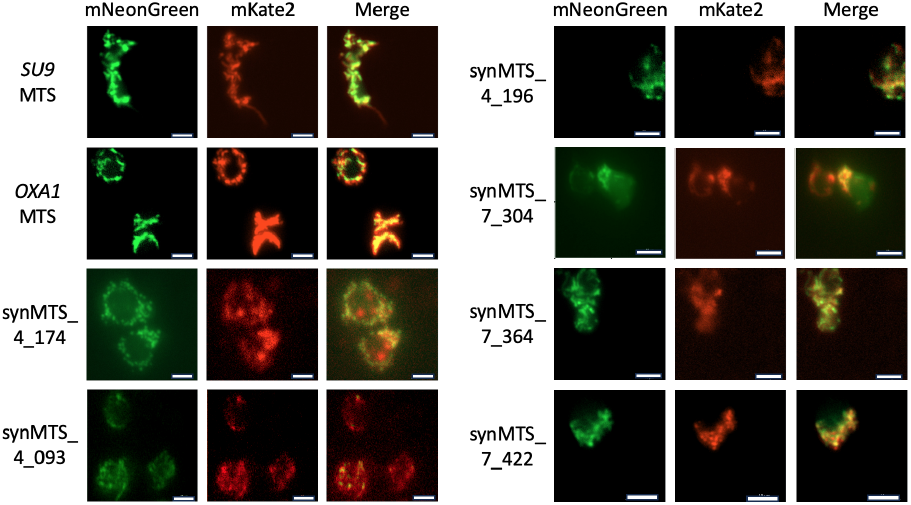
Fluorescence-based localisation assay performed in HAP1 human cells. The mNeonGreen was associated with the *COX8* MTS and used as a positive control of mitochondrial localisation while the candidate MTS/synMTS were addressing the mKate2. Scale bar = 10 *µ*m

## Discussion

In the present study, we demonstrated the possibility of creating passenger protein-specific synthetic MTS inspired from nature, which are entirely new, by using only four simple rules that capture known characteristics of MTS (see Table 1). Particularly, our aim was to generate sequences able to address artificially nucleus-relocated OXPHOS proteins (Cox2p-W56R, Atp8p, and Atp9p) in yeast to shed light on the determinants of mitochondrial proteins import.

These mtDNA-encoded proteins are usually highly hydrophobic and extremely difficult to express from the nucleus as they are highly prone to cytoplasmic aggregation, incorrect addressing or impaired mitochondrial import (37; 4). An alternative approach to improving protein addressing is to engineer their protein sequence to artificially decrease hydrophobicity. Prior reports demonstrated the benefit of adjusting the amino-acid composition on the mitochondrial import and maturation (37; 4; 15). However, a substantial fraction of these proteins still remains unprocessed in the cytoplasm, stressing the need for enhanced strategies to optimise mitochondrial uptake through engineered MTS. Therefore, introducing a straightforward approach to improve the identification of determinants for the mitochondrial import of relocated proteins would clearly help in developing efficient gene therapy strategies for mitochondrial diseases.

Due to the lack of MTS inter-compatibility (8; 15), the use of data sets of natural MTS is limited, so large data sets of *in vivo* characterised allotopic protein-specific synthetic presequences must be generated. To address this, we introduce a simple model able to generate highly functional synthetic MTS using very simple design parameters to keep a tight control on the output sequences. We implemented nine different methods, created more than 5,000 synthetic MTS *in silico* (Figure S1(A)) and benchmarked them using two software (MitoFates and DeepMito). Two promising methods (4 and 7), characterised by the most stringent rules, were identified (Figure S1(B)). The most beneficial trait identified imposed that the sequences carry a regular pattern of arginine and lysine residues, which is essential for building both amphipathic motifs and accumulated positive net charges. This is in line with the results previously published on artificial MTS (1; 33). These parameters are known to be essential for the recognition of the mitochondrial protein precursor by the mitochondrial translocases of both the outer and inner membranes (12). Using DeepMito, we obtained prediction scores indicating the high probability of passenger-protein specific synergies within our data sets (See Figure S1(C), Figure S7, and Table S11). By performing a series of *in vivo* tests in yeast on a selected subset of 50 synMTS, we not only proved their high performance but alos successfully demonstrated the existence of passenger protein-specific signatures.

In this study, we emphasize that to identify the determinants of nucleus-relocated proteins’ mitochondrial import, assays must focus on directly assessing the targeting of the proteins of interest to mitochondria (i.e. spot-assays or OCR quantification). While indirect assays can be informative (i.e. addressing a fluorescent reporter to mitochondria), it cannot be used to infer their efficacy to address allotopic proteins. Here, for each of the allotopic proteins studied, we only tested 20 out of the 1,200 generated presequences. Despite using such a small number, we could still identify highly functional sequences with clear instances of passenger protein-specific synergies, confirming the suitability of our procedure. Even though, we cannot yet precisely shed light on the amino acid composition explaining these synergies, we confirmed that: i) a regular pattern of hydrophilic residues is mandatory in defining MTS functionality; and ii) the amino acids composition in between hydrophilic residues defines the passenger protein specificity. These results highlight the uniqueness of MTS-passenger protein synergy. By further expanding our set of functionally validated synMTS, we will be able to identify the determinants that confer the ability to import nucleus-relocated OXPHOS subunits.

To conclude, the preliminary results obtained on the capability of our candidate synMTS to address the mKate2 fluorescent reporter to mitochondria in human cells are really encouraging. Subsequent investigations are required to help shedding light on the mechanisms of mitochondrial protein import and take us one step closer towards improving gene therapy-based treatment for mitochondrial diseases.

## Supplementary Data statement

Supplementary data are available online.

## Acknowledgements

The authors thank the developers of DeepMito, for promptly answering our questions about their system. The authors thank Dr. Niemi for sharing the COX4 deletion strain with us. The authors thank Pr. Herrmann for sharing the IQ-Compete system with us.

## Author Contributions Statement

K.G.: Conceptualisation, Methodology, Validation, Formal analysis, Investigation, Data Curation, Writing - Original Draft, Writing - Review & Editing, Visualisation, Supervision, Project Administration. R.W.: Conceptualisation, Methodology, Validation, Formal analysis, Software, Investigation, Data Curation, Writing - Original Draft, Writing - Review & Editing, Visualisation. J.S.J: Validation, Writing - Review & Editing. D.T.T.: Resources, Writing - Review & Editing. Y.C.: Resources, Writing - Review & Editing, Supervision, Funding Acquisition.

