## Supplementary for "TargetMITO: A rule-based model for generating highly functional synthetic mitochondrial targeting sequence in yeast"

*TBA*, YEAR, pp. 1–6
TBA
TBA
**Paper**

### TargetMITO: A rule-based model for generating highly functional synthetic mitochondrial targeting sequence in yeast

Kewin Gombeau,[1,*] Raymond Wan,[1,*] Joshua S. James,[1] Déborah Tribouillard-Tanvier[2] and Yizhi Cai[1,*]

[1]Manchester Institute of Biotechnology, University of Manchester, 131 Princess Street, M1 7DN, Manchester, United Kingdom
[2]CNRS, IBGC, UMR 5095, University of Bordeaux, F-33000 Bordeaux, France

#### Abstract

**Mitochondria are essential organelles containing their own genomes, encoding a few proteins essential for energy production. Most of the mitochondrial proteins are nucleus-encoded, translated as precursors in the cytoplasm, with a large fraction of these precursors properly addressed by an N-terminal mitochondrial targeting sequence (MTS). These MTS share common features but no consensus sequence can explain their functionality nor the precursors-specific determinants of mitochondrial import. To decipher this mechanism, we created a simple computational model to generate highly functional synthetic MTS while maintaining a tight control on the design parameters. Using the budding yeast, we demonstrated the presence of precursors-specific signatures in addressing artificially nucleus-relocated OXPHOS proteins. We also show the ability of six promising candidate synthetic MTS to address a fluorescent reporter to human mitochondria cells. Our research work confirms the uniqueness of the MTS-passenger protein synergy and takes us one step closer towards improving gene therapy-based treatment of mitochondrial diseases.**

| Amino acid | Abbreviation | Hydrophilicity |
|---|---|---|
| alanine | A | -0.5 |
| cysteine | C | -1 |
| aspartate | D | 3 |
| glutamate | E | 3 |
| phenylalanine | F | -2.5 |
| glycine | G | 0 |
| histidine | H | -0.5 |
| isoleucine | I | -1.8 |
| lysine | K | 3 |
| leucine | L | -1.8 |
| methionine | M | -1.3 |
| asparagine | N | 0.2 |
| proline | P | 0 |
| glutamine | Q | 0.2 |
| arginine | R | 3 |
| serine | S | 0.3 |
| threonine | T | -0.4 |
| valine | V | -1.5 |
| tryptophan | W | -3.4 |
| tyrosine | Y | -2.3 |

**Table S9** Amino acid hydrophilicity scale for the 20 canonical amino acids (1).

| Method | Regular expression |
|---|---|
| 1 | $\mathsf{M}(\mathcal{A})^{+}$ |
| 2 | $\mathsf{M}\mathcal{A}^{+}(\mathsf{R}\mathcal{A}\mathsf{Y}^{\{0,1\}})$ |
| 3 | $\mathsf{M}\mathcal{A}^{*}(\mathcal{H}_1^{+}\mathcal{A}^{*})^{+}(\mathsf{R}\mathcal{A}\mathsf{Y}^{\{0,1\}})$ |
| 4 | $\mathsf{M}\mathcal{O}^{\{2,4\}}(\mathcal{H}_1\,\mathcal{O}^{\{2,4\}})^{+}(\mathsf{R}\mathcal{A}\mathsf{Y}^{\{0,1\}})$ |
| 5 | $\mathsf{M}\mathcal{O}^{\{2,4\}}(\mathcal{H}_2\,\mathcal{O}^{\{2,4\}})^{+}(\mathsf{R}\mathcal{A}\mathsf{Y}^{\{0,1\}})$ |
| 6 | $\mathsf{M}\mathcal{O}^{\{2,4\}}(\mathcal{H}\,\mathcal{O}^{\{2,4\}})^{+}(\mathsf{R}\mathcal{A}\mathsf{Y}^{\{0,1\}})$ |
| 7 | $\mathsf{M}\mathcal{O}^{\{2,4\}}(\mathcal{H}_1^{\{1,2\}}\mathcal{O}^{\{2,4\}})^{+}(\mathsf{R}\mathcal{A}\mathsf{Y}^{\{0,1\}})$ |
| 8 | $\mathsf{M}\mathcal{O}^{\{2,4\}}(\mathcal{H}_2\,\mathcal{H}_1^{\{0,1\}}\mathcal{O}^{\{2,4\}})^{+}(\mathsf{R}\mathcal{A}\mathsf{Y}^{\{0,1\}})$ |
| 9 | $\mathsf{M}\mathcal{O}^{\{2,4\}}(\mathcal{H}\,\mathcal{H}_1^{\{0,1\}}\mathcal{O}^{\{2,4\}})^{+}(\mathsf{R}\mathcal{A}\mathsf{Y}^{\{0,1\}})$ |

**Table S10** Regular expressions for the 9 methods used to generate the synMTS presequences.

**Received:** Date Month Year. **Revised received:** Date Month Year. **Accepted:** Date Month Year
© The Author(s) YEAR. Copyright and licence statements to be updated by the publisher during production.

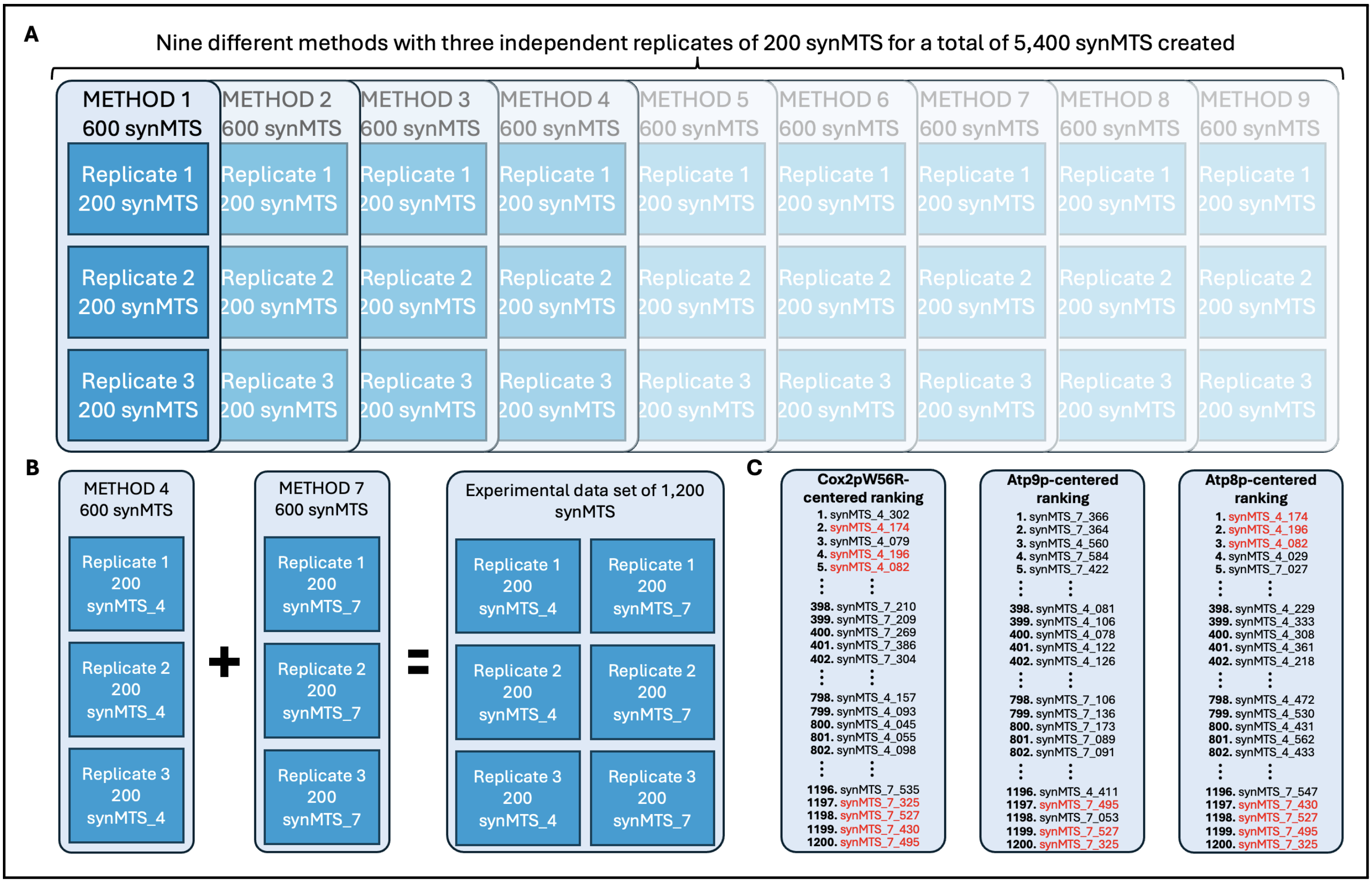


**Figure S1** Summary of the data sets created and the partitioning employed by this study. (A) Three replicates of 200 synMTS each were created for each of the 9 methods. (B) The highest scoring methods according to MitoFates were Methods 4 and 7. The synMTS of these two methods were combined into a single data set. (C) These synMTS were fused with one of three passenger proteins (Cox2p, Atp9p, and Atp8p) and ranked according to DeepMito. The synMTS highlighted in red are present in at least two data sets.

| | TPpred3/ BaCelLo | | Mitochondria compartment | | | |
|---|---|---|---|---|---|---|
| | Yes | No | OM | IM | IMS | Matrix |
| Atp8p | 1191 | 9 | 7 | 1193 | 0 | 0 |
| Atp9p | 1176 | 24 | 4 | 1196 | 0 | 0 |
| Cox2p-W56R | 1096 | 104 | 8 | 1192 | 0 | 0 |
| Hac1 | 361 | 839 | 111 | 372 | 348 | 369 |

**Table S11** Summary of the DeepMito analyses performed on the merged data sets from Methods 4 and 7 for each of the four proteins. Mitochondria localisation is reported by TPpred3 and BaCelLo as a "Yes" or a "No". Regarding the sub-compartment localisation, DeepMito calculates a probability score for each compartment and returns the highest score and the corresponding sub-compartment. The abbreviations for the mitochondrial compartments listed in the table are: OM = outer membrane, IM = inner membrane, and IMS = intermembrane space.

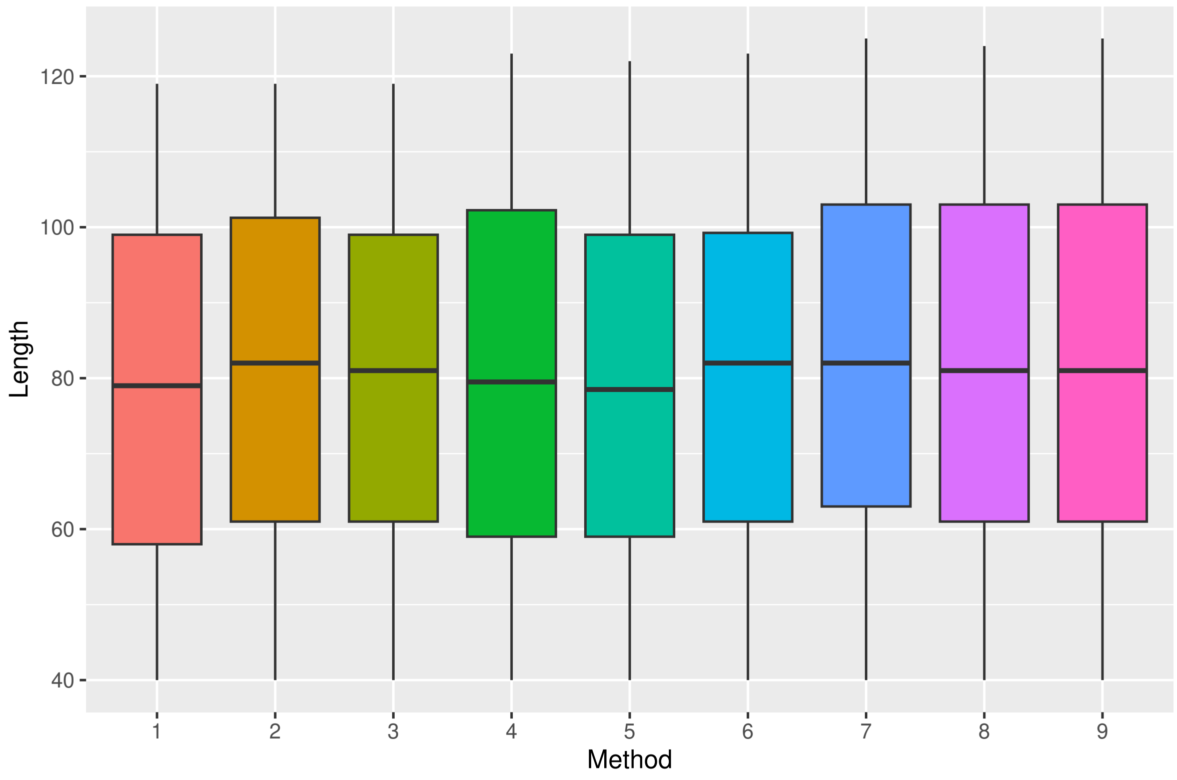


**Figure S2** Distribution of synMTS lengths for the 9 different methods.

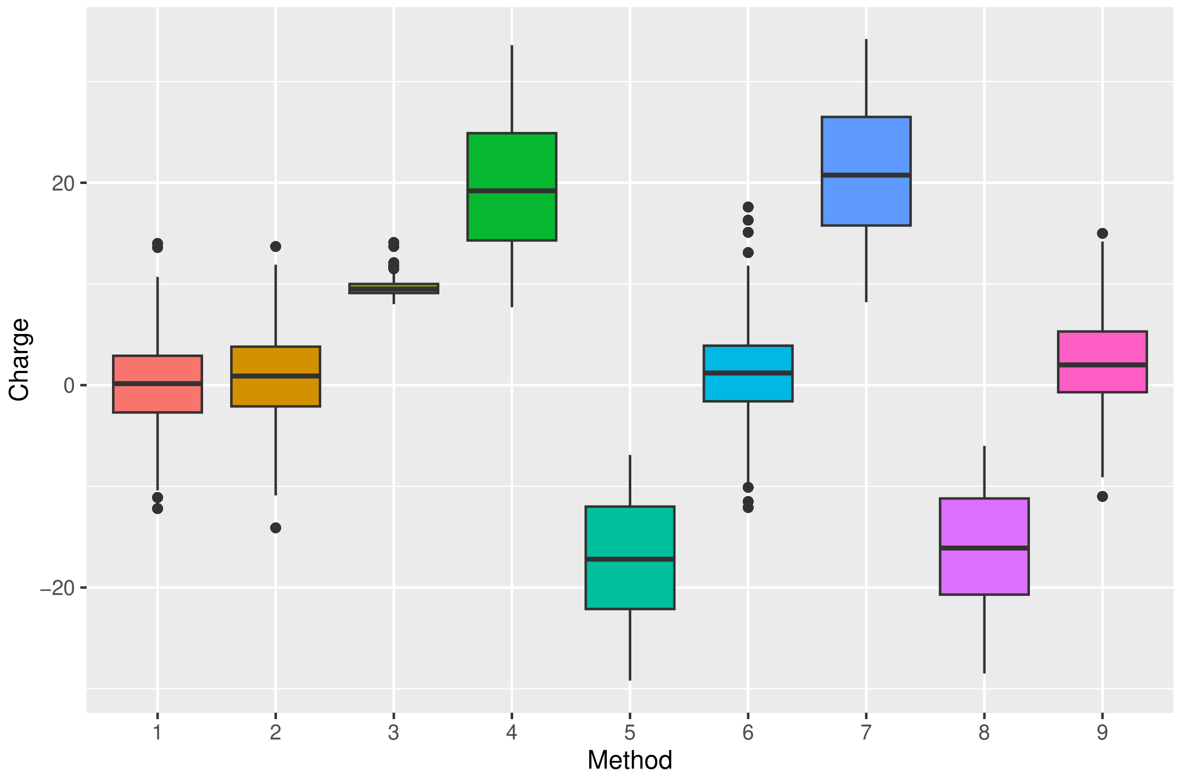


**Figure S3** Distribution of net charges of the synMTS for the 9 different methods.

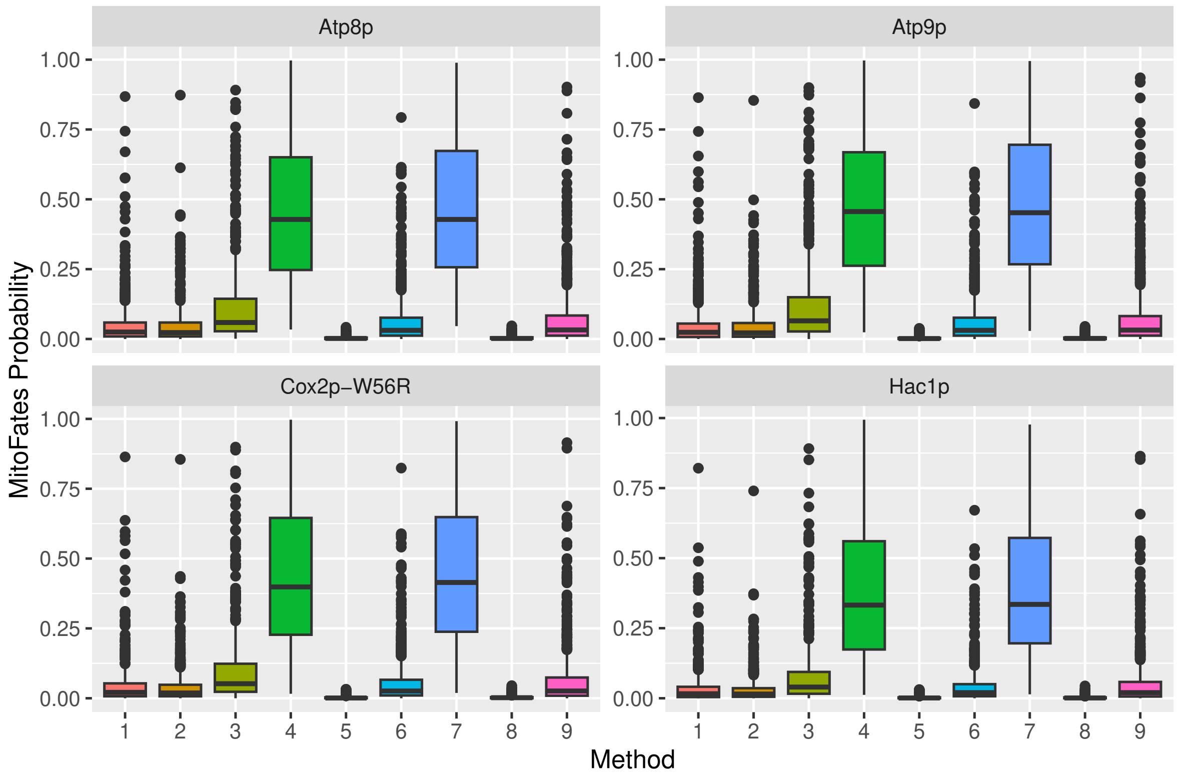


**Figure S4** Box plots of the distribution of the MitoFates presequence probability scores obtained for each method. Values have been separated according to the passenger protein appended on to the MTS sequence.

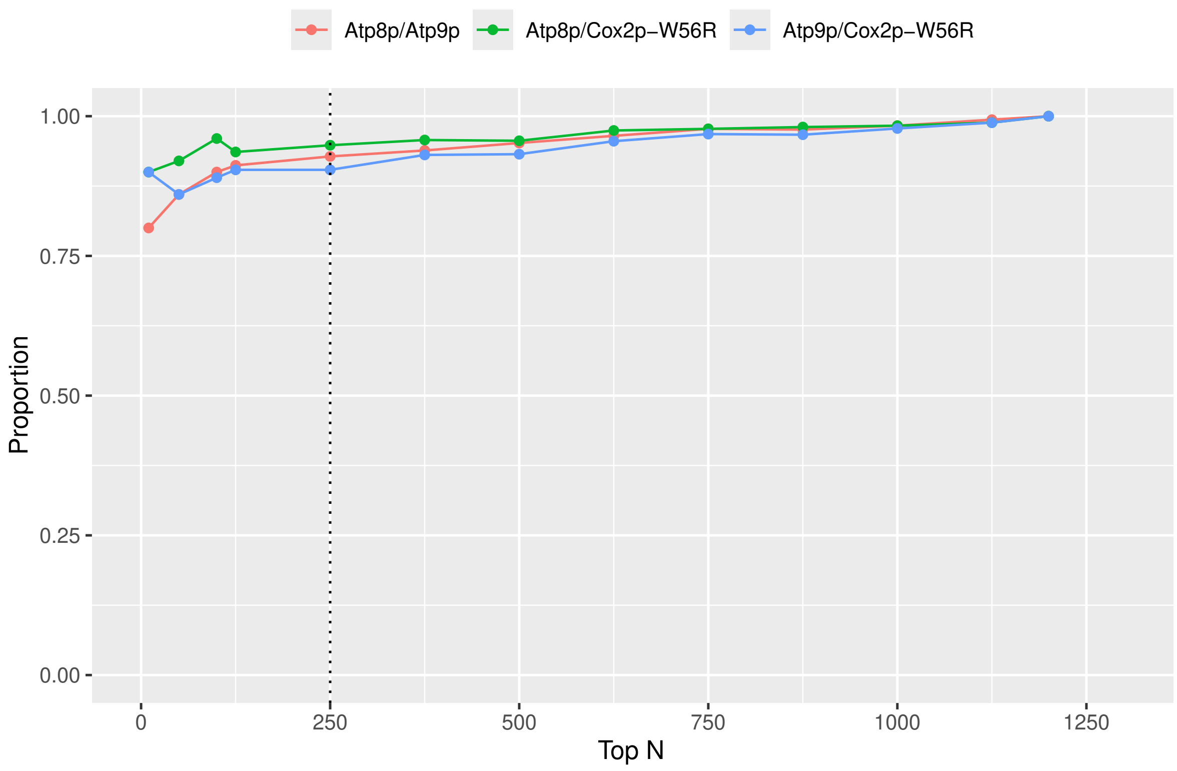


**Figure S5** Intersection of the top N MitoFates presequence probability scores for the merged data sets from Methods 4 and 7 with either Cox2p-W56R, Atp9p or Atp8p appended. The dotted red vertical line indicates the proportion of overlapping scores when only the top 250 synMTS-passenger proteins are considered.

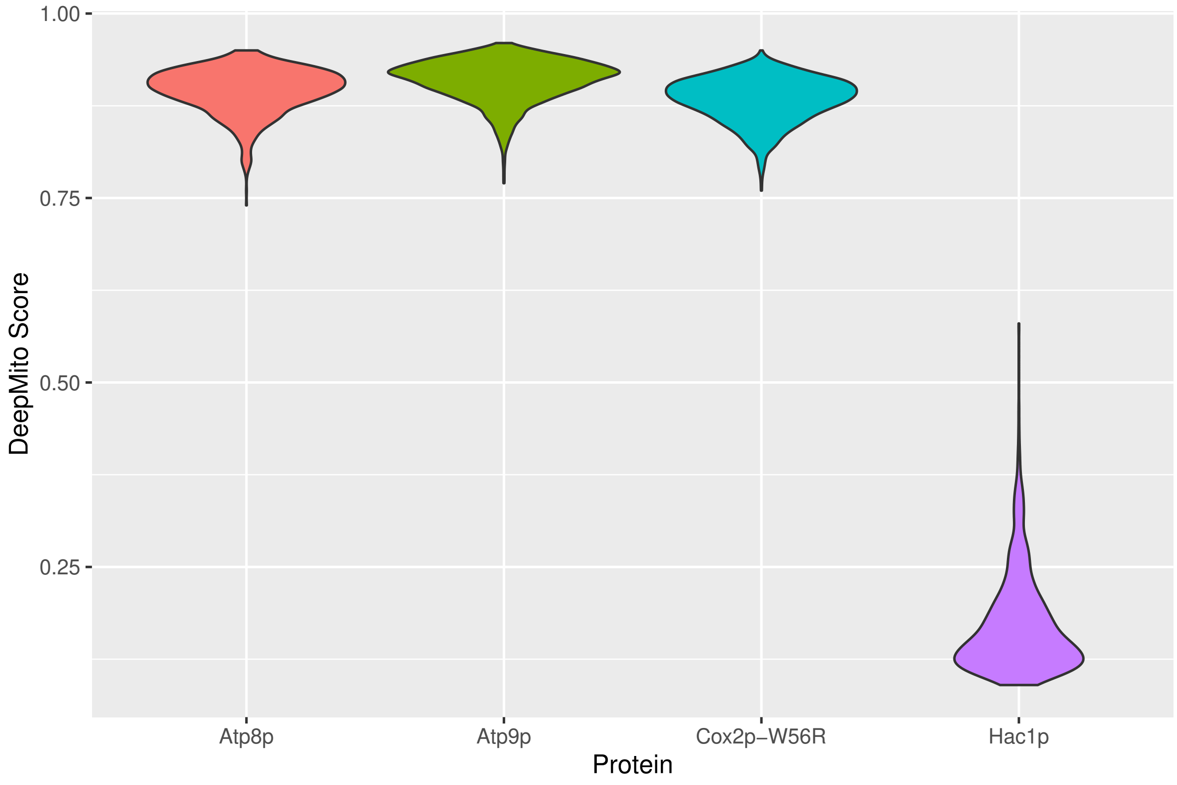


**Figure S6** Violin plots presenting the DeepMito sub-compartment localisation scores for the merged data sets from Methods 4 and 7 for each of the four proteins.

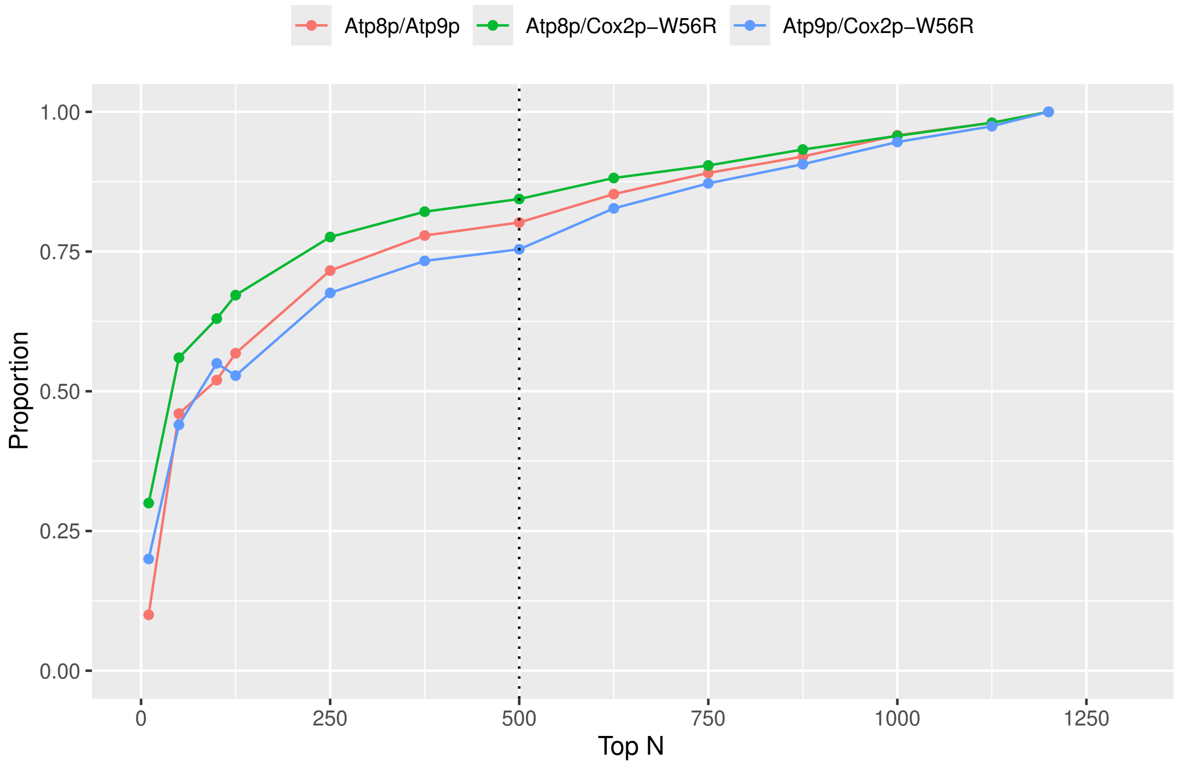


**Figure S7** Pairwise comparison of synMTS-passenger protein pairs, which shows the proportion of overlap between the two lists of DeepMito scores.
